# X-MOL: large-scale pre-training for molecular understanding and diverse molecular analysis

**DOI:** 10.1101/2020.12.23.424259

**Authors:** Dongyu Xue, Han Zhang, Dongling Xiao, Yukang Gong, Guohui Chuai, Yu Sun, Hao Tian, Hua Wu, Yukun Li, Qi Liu

## Abstract

In silico modelling and analysis of small molecules substantially accelerates the process of drug development. Representing and understanding molecules is the fundamental step for various in silico molecular analysis tasks. Traditionally, these molecular analysis tasks have been investigated individually and separately. In this study, we presented X-MOL, which applies large-scale pre-training technology on 1.1 billion molecules for molecular understanding and representation, and then, carefully designed fine-tuning was performed to accommodate diverse downstream molecular analysis tasks, including molecular property prediction, chemical reaction analysis, drug-drug interaction prediction, de novo generation of molecules and molecule optimization. As a result, X-MOL was proven to achieve state-of-the-art results on all these molecular analysis tasks with good model interpretation ability. Collectively, taking advantage of super large-scale pre-training data and super-computing power, our study practically demonstrated the utility of the idea of “mass makes miracles” in molecular representation learning and downstream in silico molecular analysis, indicating the great potential of using large-scale unlabelled data with carefully designed pre-training and fine-tuning strategies to unify existing molecular analysis tasks and substantially enhance the performance of each task.

## Main

In silico modelling and analysis of small molecules substantially accelerates the process of drug development. Representing and understanding molecules is a fundamental step towards this goal. Various molecular representations, such as molecular descriptors and fingerprints, have been proposed. Traditionally, these descriptors are designed by domain experts based on chemical and pharmaceutical knowledge to represent molecules qualitatively or quantitatively^1-4^. Various shallow learning-based machine learning models are applied to obtain quantitative structure-activity relationships (QSARs) and quantitative structure property relationships (QSPRs) to predict molecular activities and properties ^4, 5^. In recent years, with the rise of deep learning and representation learning, automatically representing and understanding molecules by learning the high-level features implied by low-level data has become an efficient method for molecular modelling, making it feasible to input the original molecules directly for subsequent molecular analysis^6-12^.

Two kinds of representations are generally adopted to represent molecules, i.e., text-based representations such as the Simplified Molecular Input Line Entry Specification (SMILES) ^13^ and graph-based representations such as 2D undirected cyclic graphs ^14-16^. SMILES is a linear representation, which is simple and straightforward and is presented as the mainstream molecular representation for deep learning modelling. In addition, taking advantage of rapidly developing natural language processing (NLP) technologies, in silico analysis of molecules based directly on text-like SMILES representations has become popular ^16, 17^. However, SMILES gains simplicity in its representations by sacrificing direct molecular structure information, which blocks its further application ^11, 18^. Due to the wide applications of SMILES and the complexity and strictness of its grammar rules, it is fundamental although challenging to understand small molecules with SMILES representations to facilitate diverse downstream in silico molecular analysis tasks, especially the task of de novo molecule generation, which has become increasingly popular in recent years ^6, 17, 19, 20^.

In this study, we explored the role of large-scale pre-training in SMILES-based molecular representation and understanding, as well as its facilitation of downstream molecular analysis tasks. Two fundamental questions related to in silico molecular modelling are investigated, i.e., (1) the feasibility of learning and understanding molecular representation well in terms of SMILES by taking advantage of massive unlabelled pre-training data and great computational power, and (2) the application of such pre-training technology to facilitate diverse downstream molecular analysis tasks. To this end, we developed a universal molecular representation learning and understanding framework, X-MOL, to facilitate comprehensive sets of molecular analysis tasks by large-scale pre-training with 1.1 billion molecules of training data and considerable computing power under the BAIDU PaddlePaddle platform ^21^. These tasks included molecular property prediction, chemical reaction analysis, drug-drug interaction prediction, de novo molecule generation and molecule optimization. We proved that in all the tasks, X-MOL achieves generalized state-of-the-art results comparable to those of existing methods with a good model interpretation ability.

## Results

### 1. The general framework of X-MOL

The general framework of X-MOL is designed in two parts, i.e., the pre-training of X-MOL and the fine-tuning of X-MOL to accommodate different molecular analysis tasks. In the pre-training stage, we trained a universal large-scale pre-training model to understand the rules of SMILES with the support of super large-scale unlabelled molecular training datasets and super-computing powers. In the fine-tuning stage, we explored how to fine-tune X-MOL to accommodate five different tasks, including molecular property prediction, chemical reaction analysis, drug-drug interaction prediction, de novo molecule generation and molecule optimization (**Fig. 1a**). Detailed descriptions of the framework can be found in ***Materials and Methods***.

**Fig. 1.**
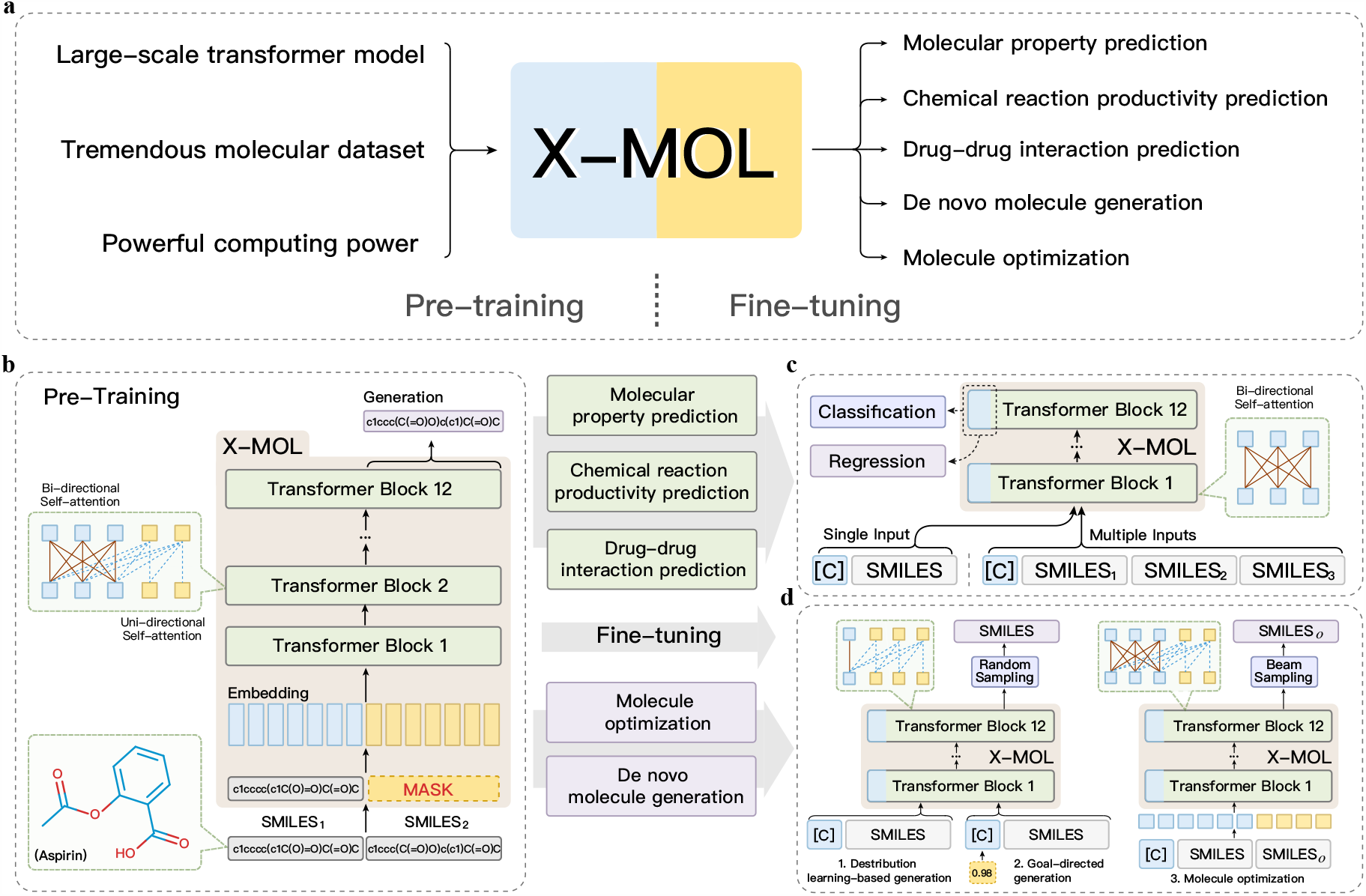
The workflow of X-MOL consists of two parts: the pre-training stage and fine-tuning stage. Pre-training is used to obtain X-MOL for SMILES representation learning and understanding, and fine-tuning is used to take advantage of X-MOL for downstream molecular analysis. **a)** The workflow of X-MOL. **b)** The shared-layer encoder-decoder structure of X-MOL in pre-training. **c)** fine-tuning X-MOL to various prediction tasks. **d)** fine-tuning X-MOL to various generation tasks.

### 2. Large-scale pre-training of X-MOL

Pre-training techniques have achieved remarkable improvements in natural language processing. Various pre-training models, such as BERT^22^, RoBERTa^23^, XLNET^24^, T5^25^ and ERNIE^26^, have been presented. The basic idea and goal of pre-training is to train the model to understand the underlying information contained in a large amount of unlabelled data and to fine-tune the model to accommodate specific downstream tasks to improve their specific performances. In our study, X-MOL adopts a well-designed pre-training strategy to learn and understand the SMILES representation efficiently. Specifically, X-MOL designs a generative model during pre-training. In this way, the model is trained to generate a valid and equivalent SMILES representation from an input SMILES representation of the same molecule. This generative training strategy ultimately results in a pre-trained model with a good understanding of the SMILES representation, and it can generate the correct SMILES of the given molecule quite well (**Fig 1a**., see ***Materials and Methods***). As a result, X-MOL builds a super large-scale pre-training model based on the Transformer^27^, which is composed of 12 encoder layers, 768-dimensional hidden units and 12 attention heads. (see ***Materials and Methods***).

### 3. Fine-tuning of X-MOL to accommodate diverse molecular analysis tasks

The universal effectiveness of X-MOL was demonstrated in five downstream molecular analysis tasks: molecular property prediction, chemical reaction analysis, drug-drug interaction prediction, de novo molecule generation and molecule optimization. According to the different task characteristics and fine-tuning strategies adopted, we divide these molecular analysis tasks into two categories, i.e., prediction tasks and generation tasks (**Fig 1b-d**). In what follows, the specific fine-tuning strategies and performances of X-MOL for these two types of tasks are presented.

#### 3.1. Fine-tuning strategies designed for two types of tasks

The detailed fine-tuning strategies designed for the two types of tasks are summarized in **Table 1**. For the prediction tasks, the specific input formats and the loss functions adopted for the different tasks are summarized. For the generation tasks, the specific generation sources and adopted sampling strategies for the different tasks are summarized (**Tab. 1**).

**Tab. 1.** Fine-tuning strategies in different downstream tasks.

| Prediction tasks |  |  |
| --- | --- | --- |
| Task | Input format | Loss function |
| Molecular property prediction (Classification) | single input | cross-entropy |
| Molecular property prediction (Regression) | single input | MSE |
| Chemical reaction productivity prediction | multiple inputs | MSE |
| Drug-drug interaction prediction | multiple inputs | cross-entropy |
| Generation tasks |  |  |
| Task | Generation source | Sampling strategy |
| Distribution learning-based generation | fixed initial symbol | random sampling |
| Goal-directed generation | unfixed initial symbol | random sampling |
| Molecule optimization | molecule | beam search |

#### 3.2. Prediction tasks

##### Molecular property prediction (classification)

Molecular property prediction, including molecular affinity and ADME/T prediction, is one of the fundamental tasks of in silico molecular analysis. In this task, we chose four benchmark classifications with few sub-datasets in MoleculeNet^28^ to test the performance of X-MOL. These four classifications are the HIV, beta-secretase (BACE), blood-brain barrier penetration (BBBP) and ClinTox datasets (see ***Materials and Methods***). The receiver operating characteristic area under the curve (ROC-AUC) was applied to evaluate the performance of the models (see ***Materials and Methods***). For the ClinToX dataset, two sub-datasets exist, and we averaged the performance of these two sub-datasets as the final performance of ClinTox (see ***Materials and Methods***). For each classification, the data were randomly split into training and testing datasets 20 times, and the average performance is reported as the final performance of X-MOL on this specific classification. A fine-tuning strategy was designed with one SMILES as the input and the probability of the sample belonging to each class as the output. The cross-entropy loss was calculated and used to update the model (see ***Materials and Methods***).

As a result, it is shown that X-MOL completely surpasses existing methods, including various classical shallow learning and deep learning models, on the four classification datasets and achieves state-of-the-art ROC-AUC values (**Fig. 2a,b,c,d**). In particular, it outperforms the existing best model significantly for the BBBP and ClinTox datasets (**Fig. 2 c,d**). It should be noted that although the graph-based molecular representation is expected to capture the molecular structure better than the SMILES representation, X-MOL still outperforms the graph-based molecular representation model GC (Graph convolutional model) ^10^, indicating that the large-scale pre-training for the SMILES representation results in a good understanding of the molecular structure and is comparable to a graph-based molecular representation.

**Fig. 2.**
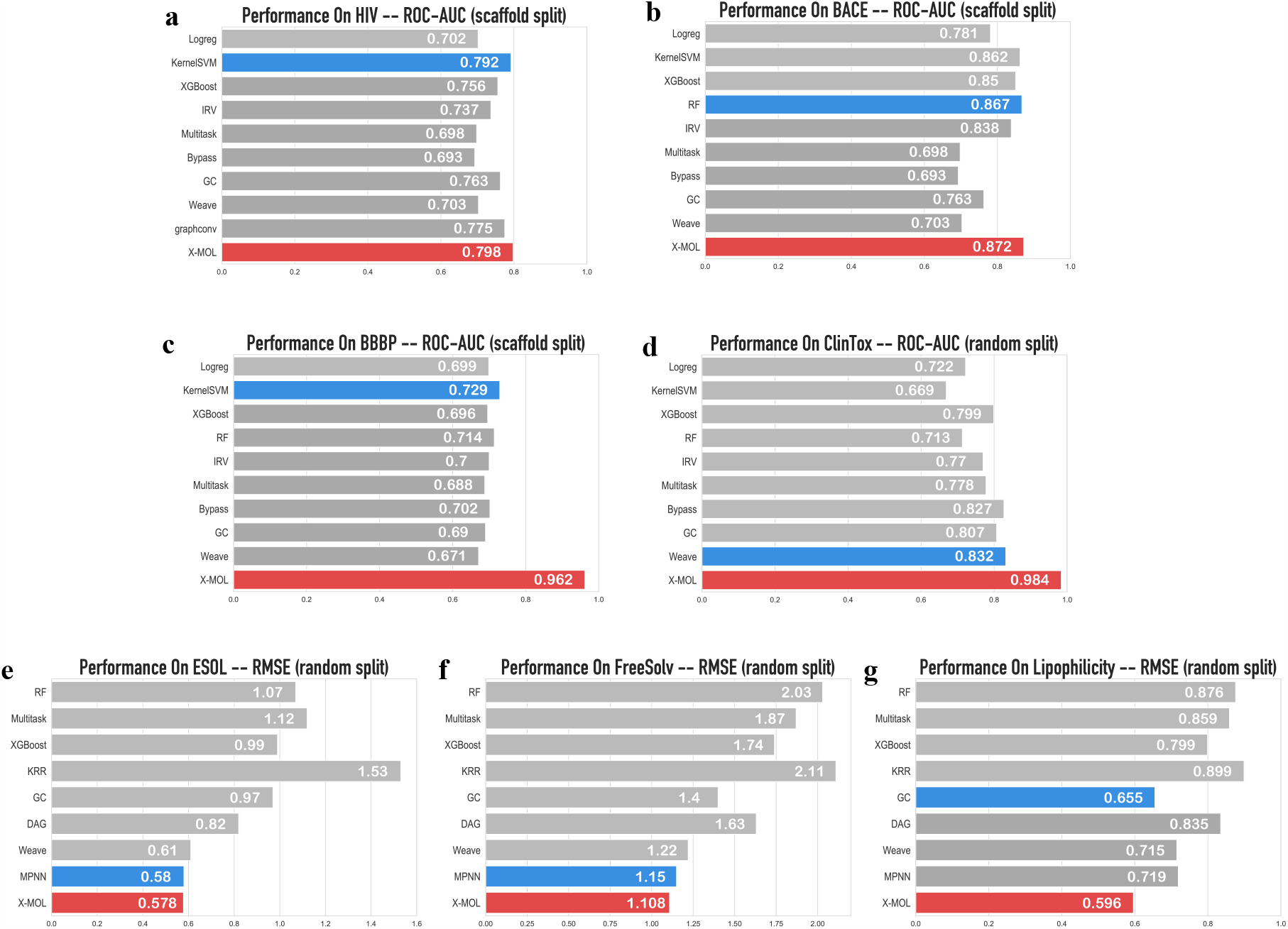
Performance comparison of X-MOL with others for molecular property prediction (Classification and Regression). **a)** Performance on the HIV dataset (Classification). **b)** Performance on the BACE dataset(Classification). **c)** Performance on the BBBP dataset(Classification). **d)** Performance on the ClinTox dataset(Classification). **e)** Performance on the ESOL dataset (Regression). **f)** Performance on the FreeSolv dataset(Regression). **g)** Performance on the Lipophilicity dataset(Regression). In the seven bar plots, the blue bar represents the best performance of the previous methods, and the red bar is the performance of X-MOL.

##### Molecular property prediction (regression)

Molecular property prediction can also be formulated as a regression task with numerical labels. In this task, three benchmark regression datasets with few sub-datasets in MoleculeNet were curated to test the performance of X-MOL for regression-based molecular property prediction. These three regressions involved the estimated solubility (ESOL), FreeSolv, and Lipophilicity datasets (see ***Materials and Methods***). The root-mean-squared error (RMSE) was applied to evaluate the performance of the models (see ***Materials and Methods***). For each regression, the data were randomly split into training and test datasets 20 times, and the average performance is reported as the final performance of X-MOL on this specific regression. After each split, a data augmentation strategy was applied to avoid over-fitting due to the insufficiency of labelled data (see ***Materials and Methods***). Similarly, a fine-tuning strategy was designed with one SMILES as the input and the predicted properties as the output. The mean squared error (MSE) loss was calculated and used to update the model (see ***Materials and Methods***).

The results show that X-MOL completely outperforms existing methods, including various classical shallow learning and deep learning models, on the three regression datasets and achieves state-of-the-art RMSE values (**Fig. 2e,f,g**).

##### Chemical reaction productivity prediction

The prediction of chemical reaction productivity is an important computational task in the field of compound production and drug synthesis. Unlike the prediction of molecular affinity or properties, the input of this task consists of several parts, including the reactants, reaction environment, catalysts and other components that participate in or affect the chemical reaction. In this task, a classical computational prediction of the reaction performance in C-N cross-coupling ^29^ was taken as the benchmark. This study included a chemical reaction productivity prediction dataset of 3956 reactions, and a random forest model (baseline) was applied for chemical reaction productivity prediction. In our study, a fine-tuning strategy was designed to take four SMILES representations as input to represent four parts of the chemical reaction, and the predicted productivity was the output. The MSE loss was calculated and used to update the model (see ***Materials and Methods***). To obtain a consistent comparison, 10-fold cross-validation was applied in our study.

As a result, X-MOL obtained an average RMSE of 0.0626, which is significantly lower than the value of 0.078 reported in the baseline (**Fig. 3a**).

**Fig. 3.**
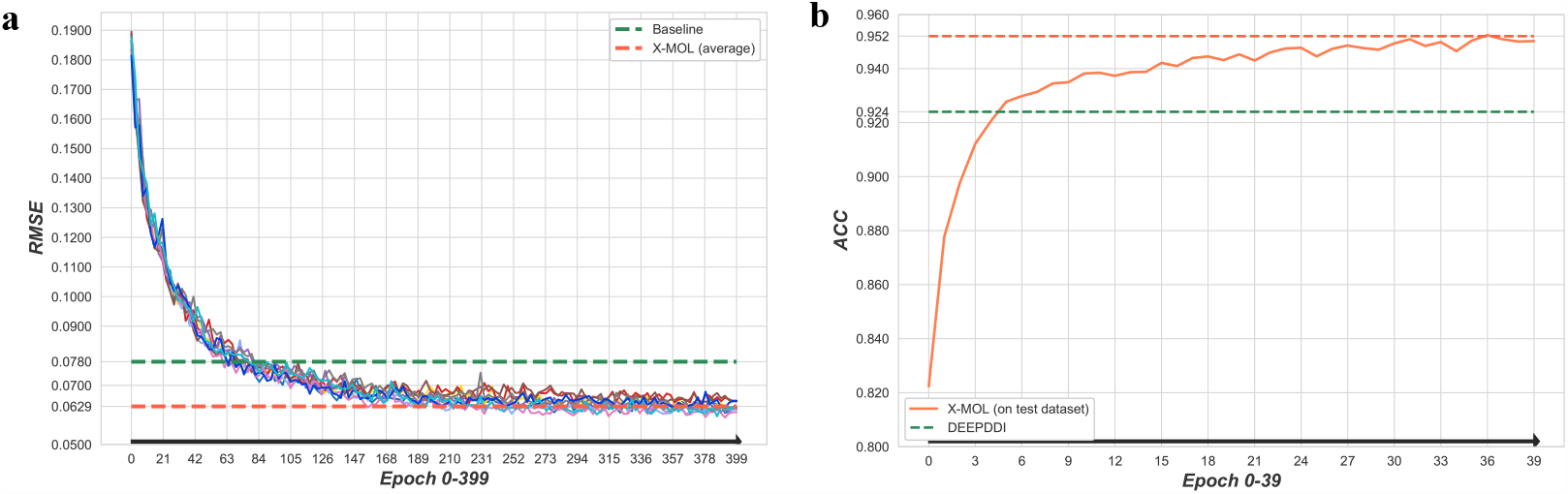
Performance comparison of X-MOL with baseline on chemical reaction productivity prediction and DDI prediction. **a)** The RMSE of X-MOL decreased during testing compared to the baseline in chemical reaction productivity prediction. **b)** The improvement of X-MOL in the DDI task during the testing.

##### Drug-drug interaction prediction

Drug-drug interactions (DDIs) may cause a series of unexpected pharmacological effects in the human body, including some unknown mechanisms of adverse drug events (ADEs); therefore, DDI prediction has become another important task related to in silico molecular analysis. In this task, a classical computational model, DeepDDI^30^, was taken as the baseline in the benchmark. This study included a DDI prediction dataset of 192284 DDIs, and a deep learning model with the designed DDI representation was presented for DDI prediction in a multi-classification way. In our study, a fine-tuning strategy was designed with two SMILESs representing two drugs as the inputs, and the probability of these drug pairs belonging to each class, for example, the increase or decrease of absorption, was the output. The cross-entropy loss was calculated and used to update the model (see ***Materials and Methods***). To obtain a consistent comparison, 10-fold cross-validation was applied in our study.

The results show that X-MOL obtained an average accuracy (ACC) of 0.952, which is higher than the value of 0.924 achieved by DeepDDI, the baseline (**Fig. 3b**).

#### 3.3 Generation tasks

The increasingly popular de novo molecule generation tasks can be divided into two categories, i.e., distribution learning (DL)-based generation and goal-directed (GD) generation^19^. In DL-based generation, the model is required to generate molecules similar to the molecules in the given training data. During the training process, the model learns the internal distribution of the training data, and then it performs sampling from the learned space to obtain new molecules. In GD generation, the model is required to generate molecules that meet certain pre-defined requirements, such as a molecular weight, quantitative estimate of drug-likeness (QED) ^31^ and LogP in a certain range. The model needs to learn the corresponding relationship between the labels of the training data and the data distribution, and then it samples and generates molecules according to the predefined goals in the process of generation.

##### Molecule generation (distribution learning-based generation)

DL-based generation is one of the most traditional generation tasks, in which the training enables the model to learn the internal distribution of the training data, and it is able to perform random sampling from the learned distribution to obtain new samples. Generally, graph-based generative models are considered to perform better than SMILES-based models in this task since a graph-based representation explicitly captures the molecular structure information. Nevertheless, in this task, we show that by taking advantage of the large-scale pre-training technology, X-MOL based on SMILES representations was able to obtain a performance in de novo molecule generation that was similar to or better than those of existing graph-based molecular representation learning models in terms of three measurements, i.e., validity, uniqueness and novelty (see ***Materials and Methods***). These existing graph-based molecule generation models include JT-VAE^32^, GCPN^33^, MRNN^34^, GraphNVP^35^, and GraphAF^36^. In particular, a fine-tuning strategy was designed with a fixed symbol as the input generation source, and a random sampling strategy was adopted for molecule generation (see ***Materials and Methods***).

As a result, X-MOL surpassed existing graph-based molecule generation methods in terms of the three measurements. In particular, X-MOL surpassed the others substantially in terms of validity (**Tab. 2a**). This result indicated that X-MOL with pre-training can learn and understand the SMILES grammar well; therefore, it can generate more valid molecules than existing methods.

**Tab. 2.**
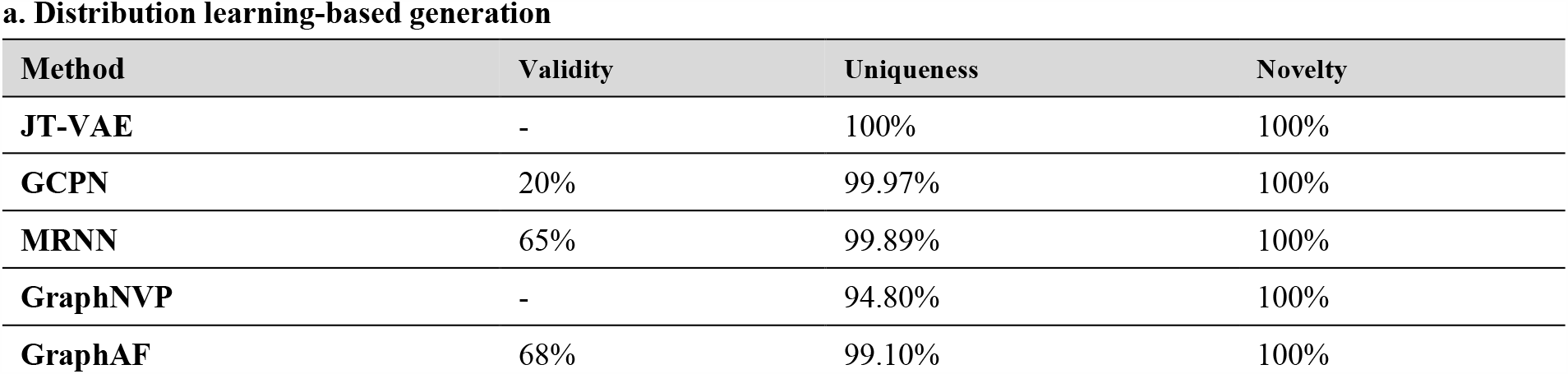
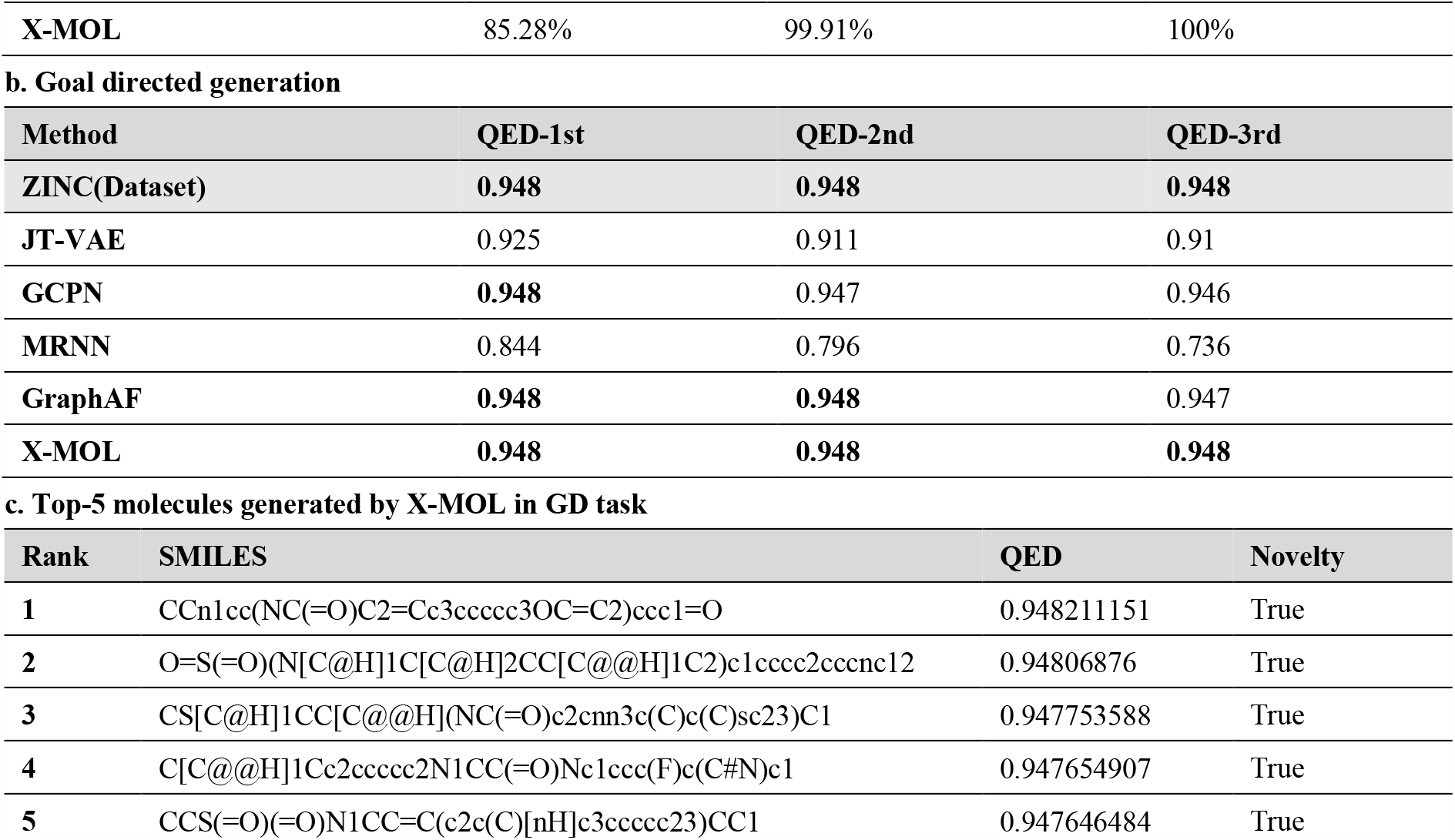
Performance comparison of X-MOL with others for de novo molecule generation. **a)** The performance comparison of X-MOL with the existing generative models in the DL task on ZINC250K dataset. **b)** The performance comparison of X-MOL with the existing generative models in the GD task on ZINC250K dataset. **c)** The top-5 molecules generated by X-MOL in GD task.

##### Molecule generation (goal-directed generation)

GD generation is another typical molecule generation task, in which a generation goal needs to be specified (see ***Materials and Methods***). Therefore, in a GD generation task, we are primarily concerned with whether the generated molecules meet the pre-defined requirements, and the performance on this task is evaluated according to the average value of the desired properties as well as the deviation from the goal value for the generated molecules. In this task, we use the zinc250k^37^ dataset as the training set to fine-tune X-MOL with the QED as the goal property; this is a classical benchmark dataset that has been widely applied in previous GD generation tasks. After the training, QED = 0.948 was set as the generation goal, since 0.948 is the maximum QED in the training set, and 10000 new molecules were generated by X-MOL for evaluation (see ***Materials and Methods***). A fine-tuning strategy was designed specifically with an unfixed symbol as the input generation source to reflect the generation goal, and a random sampling strategy was applied for GD generation (see ***Materials and Methods***).

Compared with existing graph representation-based generation models, when checking the top 3 generated molecules as in a previous study, X-MOL generated all of these molecules with the designed QED value of 0.948, outperforming graph representation-based models such as JT-VAE, GCPN, MRNN and GraphAF (**Tab. 2b**). The top-5 results are shown in **Tab. 2c**.

##### Molecule optimization

Lead compound optimization is a common step in the early stages of drug development. In this task, in silico molecule optimization was designed with an example goal to slightly modify the molecules to reach a higher QED (see ***Materials and Methods***). We investigated how large-scale pre-training and fine-tuning technology improves the final performance of molecule optimization. Therefore, X-MOL was compared with a cold-start model; i.e., this model had the same model architecture and parameters as X-MOL without pre-training or fine-tuning (see ***Materials and Methods***). The performance of molecule optimization was evaluated in terms of four measurements, i.e., goal value improvement, similarity, novelty and validity (see ***Materials and Methods***). In this task, a fine-tuning strategy was designed specifically with the molecule to be optimized as the generation source, and beam search was used for molecule optimization (see ***Materials and Methods***).

As a result, both X-MOL and the cold-start model helped to improve the target properties during molecule optimization when we compared the optimized molecules with the original ones. Nevertheless, X-MOL outperformed the cold-start model in all aspects except for the measurement of novelty (**Fig. 4c**). Specifically, we noted the following results in molecule optimization by comparing X-MOL with the cold-start model: (1) X-MOL showed a higher robustness and stability during training and testing (**Fig. 4a**). **Fig. 7b** shows that the molecules optimized by X-MOL reach at a high similarity to the original molecules shortly due to the effect of pre-training, while the similarity gradually decreased, that is desirable and completely opposite to the performance of the cold start model. (2) In terms of novelty, X-MOL showed disadvantages compared with the cold-start model, which is inevitable for generative model learning. A generative model is intended to remember the training samples during the training process; therefore, the novelty of the optimized molecules obtained by X-MOL was less than that of the cold-start model (**Fig. 4c**). (3) Since X-MOL obtains a good understanding of the grammar rules of SMILES in the pre-training stage, it can maintain a high validity during whole optimization process. Even though the cold start-model improved its validity rapidly at the beginning of optimization, a gap still existed compared with the fine-tuned X-MOL in the final convergence (**Fig. 4d**).

**Fig. 4.**
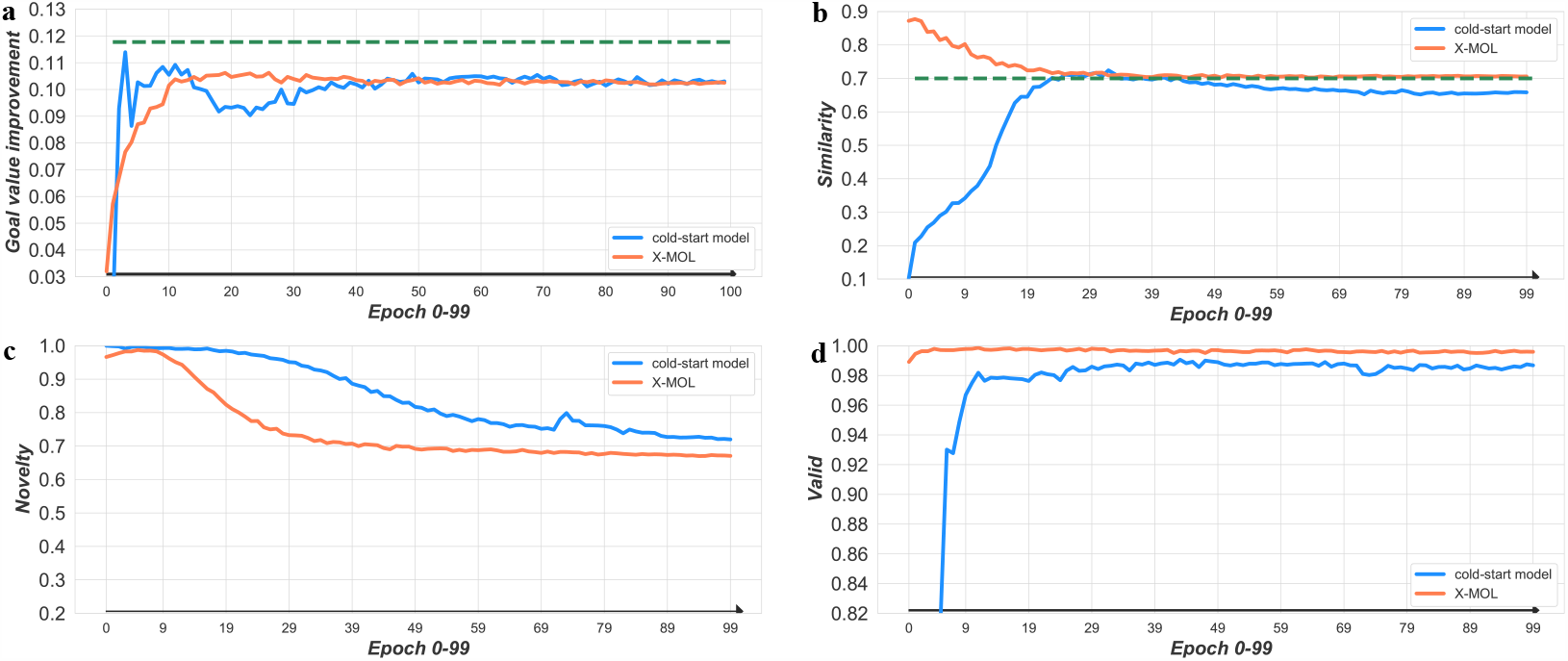
Performance comparison of X-MOL with cold-start model for molecule optimization. **a)** The average improvement in the properties of the optimized molecules compared with the input molecules. **b)** The average similarity between the optimized molecules and the input molecules. **c)** The novelty of the optimized molecules. **d)** The validity of the optimized molecules. All of the above are the processes in which the optimized molecular evaluation value changes in the optimization. Orange curve represents the fine-tuned X-MOL, blue curve indicates the cold-start model, and green line represents the average level of the training set.

#### 3.4. Visualization and interpretation of the fine-tuning for X-MOL on diverse molecular analysis tasks

Our extensive study proves that X-MOL achieves state-of-the-art performance in all molecular analysis tasks. In addition, we investigate the attention-based mechanism adopted by X-MOL in the fine-tuning stage, which allows us to intuitively check the attention of different layers on different parts of SMILES. The visualization and interpretation of such an attention-based mechanism helps us to decipher how X-MOL understands the molecules in terms of SMILES representations for different molecular analysis tasks (see ***Materials and Methods***).

For the prediction tasks, X-MOL first reconstructs the molecular structure according to the input SMILES and then infers the properties according to the molecular structure. Therefore, in the middle layer of the model, the structure of the molecules can be reconstructed accordingly, while in the later layers, the model is learned in order to extract certain high-level features of the molecular structure for specific prediction tasks. An example is shown here by investigating the molecule ***O=C(O)c1cccc(P(=O)(O)O)c1*** in the HIV dataset for molecular property prediction. The attention weights needed to learn this molecule are visualized as a heatmap, and the brightness of the colour indicates the strength of attention. The ordinate of the heatmap represents each character in the input molecule in terms of the SMILES representation, and the abscissa represents each character that is involved. Taking the character ***P*** and the last ***c1*** with the two most complex atom connection environments in the SMILES representation as examples, it is clear that in the middle layer of the model (layer 9), this method can correctly identify the environment of ***P*** and ***c1***. For ***P***, head6 recognizes the double bond connected ***O***, head7 recognizes the ***c*** connected with the benzene ring, and head11 recognizes two single bond connected ***O*** atoms. For ***c1***, head6 and head7 both pay attention to the corresponding open-loop ***c1***, while head11 recognizes a ***C*** linked to the benzene ring in addition to the open-loop ***c1***. Therefore, the two complex atom connection environments are correctly identified by X-MOL (**Fig. 5a**). Then, the model further abstracts various parts of the molecule to show their higher-level features, as seen in the later layers, and finally makes property predictions based on the degree of attention to different parts of the molecule (**Fig. 5b**). In summary, this example clearly indicates that X-MOL is able to reconstruct the non-linear molecular structure and learn its high-level features for property prediction in terms of the linear SMILES representation.

**Fig. 5.**
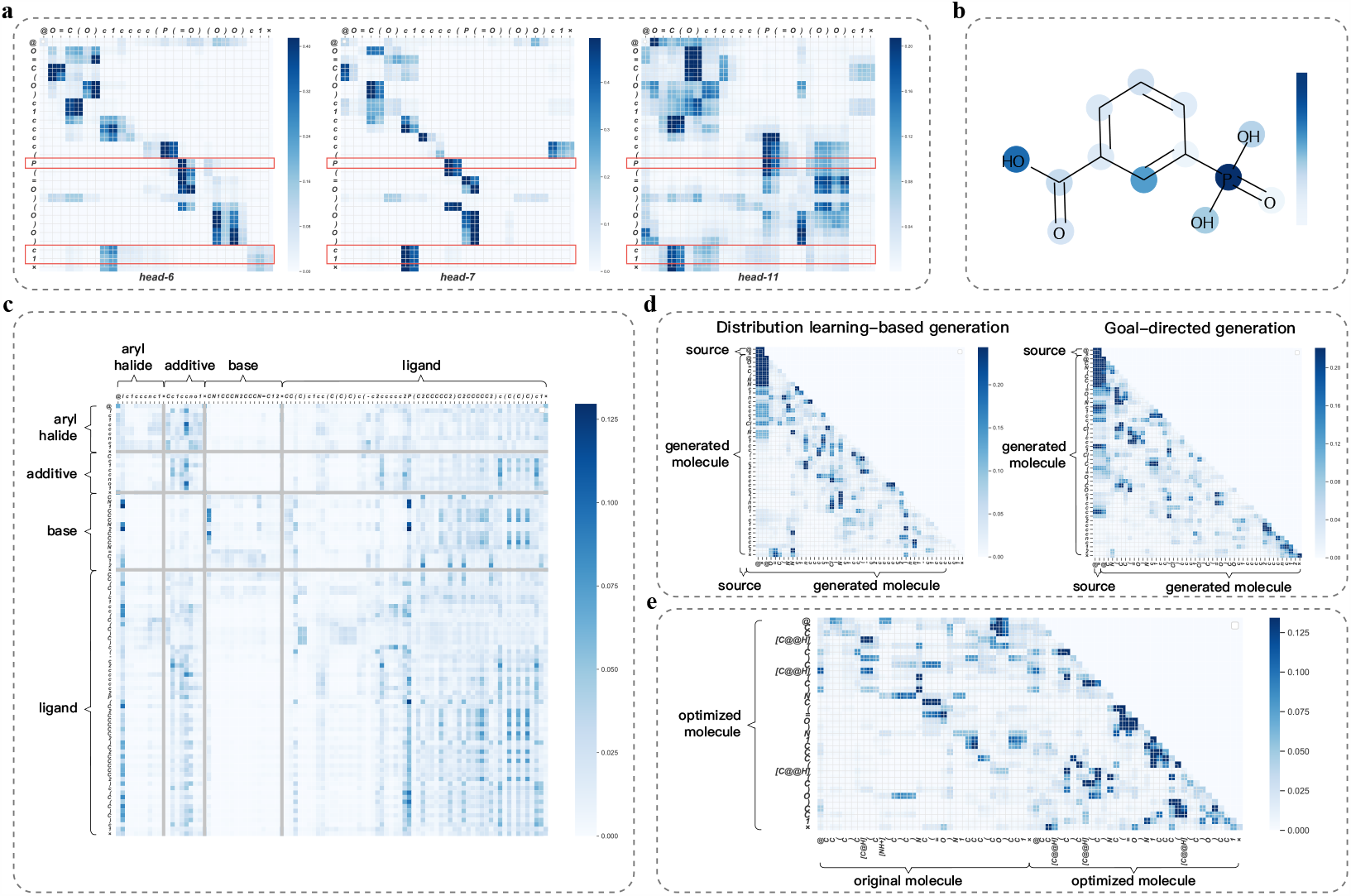
Attention visualization of X-MOL. **a)** Attention visualization of layer 9 in X-MOL on the HIV dataset for head6, head7 and head11, in molecule property prediction. **b)** The visualization of the last layer of X-MOL on the HIV dataset in molecule property prediction. **c)** Attention visualization on the reaction prediction task. **d)** Attention visualization on two de novo molecule generation tasks: distribution learning-based generation and goal-directed generation. **e)** Attention visualization on the molecule optimization task.

It should be noted that in the process of learning the features of the given molecules, the fine-tuning model is actually a process in which each part of the molecule constantly pays attention to the whole molecule and then updates itself. This is a kind of self-attention. The former example clearly indicates that self-attention is the key point in molecule affinity/property prediction. For multiple-input prediction tasks such as chemical reaction productivity prediction and DDI prediction, in addition to self-attention, attention also exists between different input components. Another demonstration of chemical reaction productivity prediction is shown in **Fig. 5c**. In this task, the reaction has four components, the ***aryl halide, additive, base*** and ***ligand***, and the ***ligand*** is the most important part because all four components have high attention to the ***ligand***. Additionally, it is clear that the ***aryl halide*** and ***base*** are less important parts—almost all of their attention is their own—while the **additive** is of concern for all components except the ***base***. This visualization of the attention is consistent with our prior chemical knowledge of this reaction, further indicating that the attention-based mechanism is efficient in understanding molecular representations under various prediction tasks.

Regarding generation tasks, the generation of new molecules is not an instantaneous process, and each SMILES is generated by one character. Therefore, in the generation process, each generated character can only focus on itself and the previously generated characters, resulting in a lower triangular attention matrix. As shown in **Fig. 5d**, the visualization of the upper-right corner of attention is generally dark for different generation tasks. However, there are specific differences among different tasks in terms of attention. In DL-based molecule generation, the model learns the generation space of the target molecules during training, and the newly generated characters pay more attention to the generated parts of themselves. In GD-based molecule generation, however, the model also needs to pay attention to the source at the end of the generation process to constrain the molecules generated following the generation goal. Moreover, in GD-based molecule generation, when the model generates new molecules, it should not only meet the grammar rules of SMILES but also meet the generation goal, which is reflected in the attention visualization as the fact that the newly generated character needs to pay attention to the source and the previously generated part simultaneously. Finally, in molecule optimization, since the model generates a new molecule with a given molecule as input, the attention in this task includes both unidirectional self-attention and encoder-decoder attention that focuses on the original input molecule. As shown in **Fig. 5e**, it is clear that the model pays more attention to the original molecule in the early stage of generation, while it gradually shifts from the original molecule to the generated part of the new molecule during optimization.

#### 3.5 SMILES is all you need

Our extensive study proved that X-MOL learned the grammar rules of SMILES through pre-training; however, we further considered whether information in addition to the SMILES representation would help to improve our model performance in prediction tasks. Here, we present ***knowledge embedding***, which is able to incorporate certain prior knowledge into the model by adding more embedding components to the input. As an exploratory study, we designed three different knowledge embedding strategies to incorporate the interpretation information related to SMILES into the model, i.e., link embedding, ring embedding and type embedding (**Fig. 6a**).

**Fig. 6.**
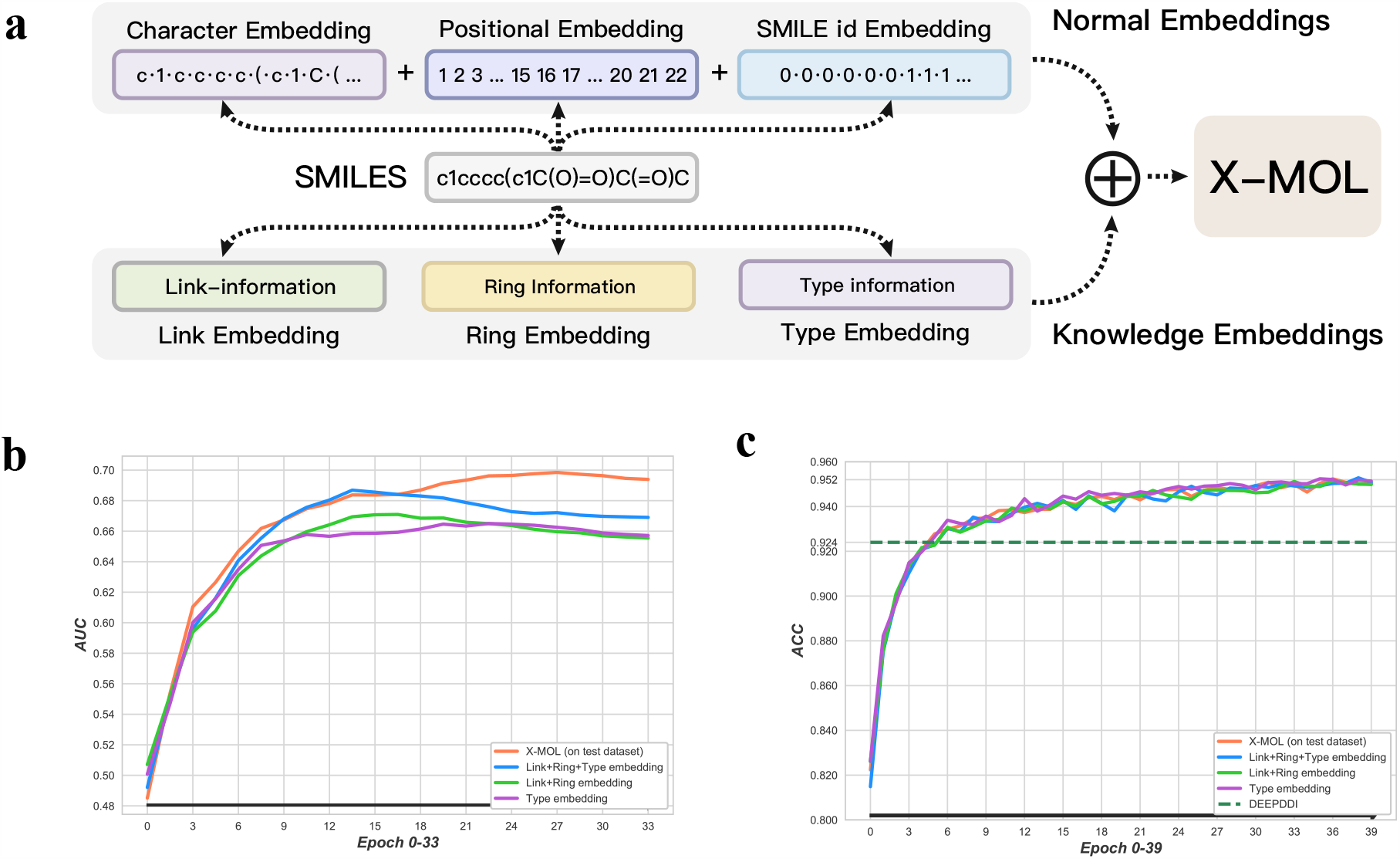
Knowledge embedding for X-MOL. **a)** The way of incorporating knowledge embedding into X-MOL. **b)** Knowledge embedding in HIV dataset for molecule property prediction. **c)** Knowledge embedding in the DDI prediction task.

##### Link embedding

Because of the grammar rules of SMILES, two adjacent atoms or functional substructures in a molecule may be far away in the SMILES representation; however, such linking information can be incorporated. To this end, we designed link embedding to incorporate the connection information of each atom, bond and symbol of the structure information in SMILES. For each character, the corresponding link embedding is the location of the previous atom to which it is connected, and the link embedding of the first atom in SMILES is represented by its own position. In this way, the linking information in SMILES is incorporated.

##### Ring embedding

SMILES uses a pair of numbers to represent a ring structure. Two identical numbers represent an open-loop atom and a closed-loop atom. In this case, the number represents the connection information (the connection between the open- and closed-loop atoms) and the ring structure. We designed ring embedding to incorporate the number pairs of the ring structure information. In ring embedding, each number indicating an open- or closed-loop atom is represented by the position of the corresponding atom. For example, a number indicating an open loop is represented by the position of the closed-loop atom, and the other characters are recorded as ***NULL***. In this way, the model can directly obtain the connection information and ring structure information represented by the ring number pair.

##### Type embedding

Because of the grammatical rules of SMILES, additional characters need to be introduced to represent the structural information of molecules when constructing SMILES, resulting in many different types of characters being used in SMILES. For example, ‘C’, ‘N’, ‘O’ and ‘[NH +]’ are used to represent atoms; ‘-’, ‘=‘ and ‘#’ are used to represent chemical bonds; and ‘(‘, ‘)’, and ‘@’, as well as paired numbers, are used to represent the molecular structure. These are completely different types of characters, and the different types of characters should be considered differently. To this end, we designed type embedding, which divides the characters into nine categories to incorporate their type information.

Finally, we selected the molecule property prediction task (HIV dataset) and the DDI prediction task as two exploratory studies. Our test results indicate that employing all the embedding strategies cannot enhance the performance of the model in the corresponding task (**Fig. 6b, c**); on the contrary, this additional information may make learning more difficult for the model. Regardless of whether the pre-training strategy is applied, these three kinds of embedding strategies may decrease the model performance to a certain degree. These results indicate that (1) the SMILES representation may contain all the information needed to obtain a good representation of the molecule, and (2) pre-training is able to extract such information and helps in learning and understanding the molecular SMILES representation well.

## Discussion

In this study, for the first time, we applied large-scale pre-training technology for molecular understanding and representation and designed X-MOL for various downstream molecular analysis tasks based on a fine-tuning strategy. We comprehensively investigated the pre-training strategy, the fine-tuning strategy, and the visualization and interpretation of the attention mechanism in X-MOL. Finally, X-MOL was proven to achieve state-of-the-art results in various downstream molecular analysis tasks, including molecular property prediction, chemical reaction analysis, drug-drug interaction prediction, de novo molecule generation and molecule optimization. These downstream molecular analysis tasks were investigated individually and separately in a previous study, while our study indicates that all of these tasks can be unified in one framework by large-scale pre-training and carefully designed fine-tuning, resulting in state-of-the-art performance that takes advantage of super large-scale unlabelled data and super-computing power. The idea of “mass makes miracles” supported by our study indicates the great potential of using large-scale unlabelled data and representation-based learning to unify existing molecular analysis tasks and to enhance each task substantially. This is of great significance for the molecular and pharmaceutical communities since the molecular space is large and rapidly increasing.

Further improvements of X-MOL are expected, including (1) better defined pre-training strategies to learn the grammar rules of SMILES; (2) the application of these pre-training strategies to other molecular representations, such as graph-based representations, and (3) fine-tuning on additional kinds of downstream molecular analysis tasks.

## Methods

### 4.1 Pre-training of X-MOL

#### The generative pre-training strategy designed for X-MOL

Pre-trained on large-scaled text corpora and finetuned on downstream tasks, self-supervised representation models like BERT, RoBERTa, ERNIE and XLNET have achieved remarkable improvements in NLP.. The basic idea of large-scale pre-training is to make the model understand the semantic information contained in large-scale unlabelled training data; then, the well-trained model can be fine-tuned to accommodate specific downstream tasks and improve the performance on these tasks. The key point of pre-training is the method of making the model understand the semantic information. For the pre-training in NLP, the masked language model (MLM) ^38^ and permuted language model (PLM) ^24^ are used to train a model to learn the semantics of a whole sentence. In our study, we design a specific generative pre-training strategy in X-MOL for the representation and understanding of SMILES.

Inspired by the characteristics of random SMILES (see **Random SMILES in Section 4.1.2**), i.e., that one molecule can have multiple valid random SMILES representations, we assume that if a model can learn the grammatical rules of SMILES well, it should achieve two goals: (1) Given a SMILES representation of a molecule, the model is able to reconstruct the corresponding structure of the molecule; (2) Given the structure of a molecule, the model is able to generate a valid SMILES representation following the grammar rules of SMILES. Based on this hypothesis, we propose a generative pre-training strategy; i.e., the pre-trained model is trained to generate another valid SMILES of the same molecule with a given SMILES as input. Once the model can perform this task, we consider that the model can not only reconstruct the molecular structure through SMILES but also generate a valid SMILES representation according to the molecular structure. In other words, the model is trained to generate valid and equivalent SMILES representation with an input SMILES representation of the same molecule.

Specifically, our generative pre-training strategy is implemented through an encoder-decoder architecture. Unlike traditional encoder-decoder architectures such as those used in neural machine translation (NMT), in X-MOL, the encoder and decoder share the parameters of a multi-layer Transformer (**Fig. 1b**) ^39^, forcing the process of encoding and decoding to be performed in the same semantic space. In X-MOL, the input random SMILES and output random SMILES are sent into the model simultaneously, and the output random SMILES is totally masked. In addition, only a unidirectional attention operation can be performed within the output random SMILES, which means that each character in the output random SMILES can pay attention only to itself and the previously generated characters.

#### Detailed technologies applied in the pre-training of X-MOL

##### Random SMILES

According to the grammar rules of SMILES, when we transform molecules into the SMILES format, we need to choose an atom as the starting point, a chain at the branch as the main chain, and the position of the open ring in the ring structure. For a specific molecule, different choices result in different SMILESs. Therefore, each SMILES accurately represents a certain molecule, but a molecule can be represented by multiple SMILESs. We denote the SMILES obtained by a depth-first algorithm as the canonical SMILES ^40^, and there is only one canonical SMILES per molecule; the SMILES obtained from a random selection of the starting site, main chain and ring opening position is called a random SMILES.

##### Data augmentation in pre-training

Due to the non-uniqueness of random SMILES, although the model can correctly reconstruct the molecular structure and return another random SMILES of the molecule, the output SMILES may be different from the predefined output random SMILES, which may confuse the model training in the design of the loss function. To solve this problem, we introduce data augmentation technology in the pre-training stage. For each molecule, we randomly generate multiple training samples, where these samples share the same input SMILES and have different outputs. During the pre-training process, these training samples with the same input should be put in the same mini-batch to guide the model in addressing such scenarios involving non-uniqueness.

##### Segmentation of SMILES

Word segmentation is a key technology in text-based NLP tasks ^41^. The method of segmentation and the level of granularity have a great impact on the task performance. Generally, word-level granularity brings more semantic information than character-level granularity. In the format of SMILES, word-level granularity corresponds to the level of the sub-structures of molecules, however, it is difficult to perform segmentation at the level of sub-structures in SMILES, since a sub-structure cannot be represented in a linear text-like format. Therefore, we perform segmentation at the character level for SMILES, where characters within a pair of square brackets and double digits with a “%” in front of them are regarded as one character. In this way, we segment all 1.1 billion SMILESs and obtain a character dictionary containing 108 characters (in addition to the 108 chemical characters, there are another 5 characters for building tasks). ***[PAD]*** is used to unify the length of the input SMILESs, ***[CLS]*** is used to record classification information or indicate the beginning of a SMILES, ***[SEP]*** is used to split multiple SMILESs, ***[MASK]*** is used for mask operations and ***[UNK]*** is used to represent unknown characters.

##### Model capacity in pre-training

With great computing power, X-MOL built a super large-scale pre-training model based on the Transformer architecture, which is composed of 12 encoder layers, 768-dimensional hidden units and 12 attention heads.

The large-scale pre-training data for X-MOL were curated as follows: We selected all the data in the ZINC15 database ^42^, with 1.1 billion molecules in total, and we used RDKit ^43^ to convert the canonical SMILESs of these molecules into different random SMILESs. In the data pre-processing stage, we used more than 1000 CPUs, combined with Hadoop technology ^44^, to process these training data at the same time.

We used BAIDU’s PaddlePaddle framework to implement the whole pre-training model and the specific downstream tasks. For each downstream task with pre-training, we used 8/16 Tesla P40 GPU (24 GB) to train the model simultaneously, and the pre-training stage took almost 4 days.

### 4.2 Fine-tuning of X-MOL

#### Categorization of fine-tuning strategies according to different molecule analysis tasks

The fine-tuning of X-MOL to accommodate different types of downstream tasks, i.e., prediction tasks and generation tasks, also plays an important role. First, in both the pre-training stage and the fine-tuning stage, we add the initial character ***[CLS]*** to the first bit of the input of the model. For different types of tasks, ***[CLS]*** plays a different role. In the prediction tasks, we use ***[CLS]*** to extract the predicted properties that we care about, while in the generation tasks, ***[CLS]*** is the initiator of the generation. The detailed fine-tuning strategies designed for the two types of tasks are summarized in **Table 1**. We explain the two types of fine-tuning strategies in detail below.

#### Fine-tuning strategies designed for the prediction tasks

In the prediction tasks, as shown in the **Figure 1c**, for single-input tasks, such as the molecular property prediction task, we directly add ***[CLS]*** and ***[SEP]*** to both sides of the molecules as the model input. For multiple-input tasks, such as chemical reaction productivity prediction and drug-drug interaction prediction, a SMILES complex is obtained by separating the multiple-input SMILESs with ***[SEP]***. Finally, ***[CLS]*** and ***[SEP]*** are added to both ends of the complex.

On the other hand, we use ***[CLS]*** to extract the features of molecules, so for the output of the model, ***[CLS]*** is all we need. Then, the features of ***[CLS]*** are connected to a fully connected network, and finally, the properties of the given molecule are output. The difference between the classification task and regression task lies in the number of neurons in the last layer of the fully connected network and their activation functions (**Fig. 1c**).

#### Fine-tuning strategies designed for the generation tasks

For the generation tasks, the architecture is quite similar to that of the pre-training model. For the DL-based generation task, the model learns the chemical distribution within the data and generates new molecules without input. In contrast to the pre-training model, we remove the reference SMILES in the input and only send ***[CLS]*** and ***[SEP]*** to the model as the input to generate new molecules. In this model, ***[CLS]*** and ***[SEP]*** are regarded as the starting characters of the training process. For the GD task, the overall structure is similar to that of the DL-based generation tasks, while the GD task needs the input of the generation goal. Therefore, we modify the input ***[CLS]*** and combine the modified ***[CLS]*** with the unmodified ***[SEP]*** as the new input of the generation model.

In addition, the sampling strategies applied for different generation tasks are different. DL-based de novo molecule generation and GD de novo molecule generation have multiple generation results, while the molecule optimization task generates only one result for a given input molecule. Therefore, random sampling is applied for de novo molecule generation, and beam search (here, we use beam-size=4) was applied for molecule optimization (**Fig. 1d**).

#### Detailed technologies applied in the fine-tuning of X-MOL

##### Repeated training

During the fine-tuning tasks, it was found that if the amount of training data is insufficient, the overall data distribution is unstable. When the data are randomly divided into a training set and a test set, the training set and test set may be too similar or too different, which will substantially affect the performance of the model on the test set. To make the model performance more reliable, a repeated training strategy was applied. A complete training process includes data splitting and model training, we repeated the training 20 times. In each training process, the data were randomly split into a training test set (or training, validation and test set), and then X-MOL was re-trained on the training set and tested on the test set. Finally, the average performance of the 20 models obtained from 20 repeated trainings is reported as the final performance of X-MOL.

##### Data augmentation

We applied data augmentation in the prediction tasks. In most molecular datasets, the molecules are stored in the canonical SMILES format. After randomly splitting the dataset into training, validation and test sets, we augmented the training set by adding random SMILES which is transformed from canonical SMILES with the validation and test sets unchanged.

##### Design details of the fine-tuning of X-MOL for different molecular analysis tasks Molecular property prediction (classification)

We choose four classification datasets with few sub-datasets in MoleculeNet to test the performance of X-MOL on classification tasks. These four datasets are HIV, BACE, BBBP and ClinTox.

The **HIV** dataset was introduced by the Drug Therapeutics Program (DTP) AIDS Antiviral Screen, which tested the ability of over 40,000 compounds to inhibit HIV replication. The screening results were evaluated and placed into three categories: confirmed inactive (CI), confirmed active (CA) and confirmed moderately active (CM). We combined the last two labels, making this a classification task between inactive (CI) and active (CA and CM).

The **BACE** dataset provides quantitative (IC50) and qualitative (binary label) binding results for a set of inhibitors of human β-secretase 1 (BACE-1). All the data are experimental values reported in the scientific literature over the past decade, and some have detailed crystal structures available. A collection of 1522 compounds with their 2D structures and properties are provided.

The **BBBP** dataset was extracted from a study on the modelling and prediction of barrier permeability. As a membrane separating the circulating extracellular fluid in the blood and brain, the blood-brain barrier blocks most drugs, hormones and neurotransmitters. Thus, penetration of the barrier is a long-standing issue in the development of drugs targeting the central nervous system. This dataset includes binary labels for over 2000 compounds concerning their permeability properties. The **ClinTox** dataset compares drugs approved by the Food and Drug Administration (FDA) and drugs that have failed clinical trials for toxicity reasons. The dataset includes two classification tasks for 1491 drug compounds with known chemical structures: (1) clinical trial toxicity (or absence of toxicity) and (2) FDA approval status. The list of FDA-approved drugs was compiled from the SWEETLEAD database, and the list of drugs that failed clinical trials for toxicity reasons was compiled from the Aggregate Analysis of ClinicalTrials.gov (AACT) database.

The amounts of data in these four datasets are shown in **Table 3**. We use the ROC-AUC to evaluate the performance of the model according to the benchmark in MoleculeNet. Repeated training was applied for these four datasets. There are two sub-datasets in ClinTox, and we averaged the performance of the two sub-datasets as the final result for the ClinTox dataset.

**Tab. 3.** The data amount for different datasets in the molecule property prediction task.

| Dataset name | Data amount | Sub-dataset number | Task type |
| --- | --- | --- | --- |
| **HIV** | 41127 | 1 | classification |
| **BACE** | 1513 | 1 | classification |
| **BBBP** | 2039 | 1 | classification |
| **ClinTox** | 1484 | 2 | classification |
| **ESOL** | 1128 | 1 | regression |
| **FreeSolv** | 642 | 1 | regression |
| **Lipophilicity** | 4200 | 1 | regression |

##### Molecular property prediction (regression)

All three regression datasets with few sub-datasets were selected from MoleculeNet, including ESOL, FreeSolv and Lipophilicity.

**ESOL** is a standard regression dataset containing structures and water solubility data for 1128 compounds. This dataset is widely used to validate machine learning models for estimating solubility directly from molecular structures.

The **Free Solvation** Database, FreeSolv (SAMPL), provides the experimental and calculated hydration free energy of small molecules in water. The experimental values are included in the benchmark collection, and the measured solvation energies (unit: kcal/mol) of the compounds are used as the labels in this task. **Lipophilicity** is a dataset curated from the ChEMBL database containing experimental results on the octanol/water distribution coefficient (logD at pH=7.4). Due to the importance of lipophilicity in membrane permeability and solubility, this task is of high importance in drug development.

Compared with that of the four classification datasets, the data volume of the three regression datasets is smaller (**Tab. 3**); therefore, in addition to repeated training, data argumentation is applied here. We use the RMSE to evaluate the performance of the model according to the benchmark in MoleculeNet.

##### Chemical reaction productivity prediction

The input of this task consisted of four parts, an aryl halide, additive, base and ligand, and the output was the productivity of the chemical reaction. The data provided by the reference study contain a total of 3956 reactions and were not previously split; we split the dataset into a training set and a test set according to the ratio of 7:3 in the original study. The RMSE was taken as the evaluation measurement, as in the reference study.

##### Drug-drug interaction prediction

The dataset provided by the reference study contains 192284 DDI pairs, and they were split into training, validation and test sets. There are 86 types of DDIs in the DeepDDI data, so the DDI prediction task was formulated as a multi-classification task with two input parts. The ACC was taken as the evaluation measurement for DeepDDI.

##### Molecule generation (distribution learning-based generation)

The ZINC250K dataset was used in this task as the benchmark dataset containing 249456 molecules. The performance was evaluated in terms of three measurements, i.e., validity, uniqueness, and novelty. Validity represents the proportion of generated molecules that meet the rules of the SMILES grammar. Uniqueness represents the proportion of non-repetitive samples among the generated molecules. Novelty represents the proportion of new samples among the generated molecules that are absent from the training data, indicating the ability of the model to generate novel molecules.

##### Molecule generation (goal-directed generation)

The ZINC250K dataset was used in this task as the benchmark dataset, containing 249456 molecules. The QED value was set as the generation goal, which can be calculated by RDKit.

In the implementation of the model, our design was based on class-oriented molecule generation, where an embedding layer is constructed to convert a class into a feature vector represented by floating-point numbers. Generally, ***[CLS]*** (the starting character of generation, also called ***[BOS]*** and ***[start]*** in other works) is used to represent different classes in class-oriented generation. After the transformation of the embedding layer, ***[CLS]*** becomes a feature vector composed of floating-point numbers. Then, the model generates different categories of molecules according to different ***[CLS]*** eigenvectors. Value-oriented generation is designed by modifying the eigenvector of the original ***[CLS]***. We multiply the value of the generation goal with the ***[CLS]*** eigenvector obtained after pre-training to obtain the new eigenvector of ***[CLS]*** in value-oriented GD tasks, incorporating the goal value by passing through the embedding layer.

##### Molecule optimization

The ZINC250K dataset was used in this task as the benchmark dataset, containing 249456 molecules. In this task, the fine-tuning model was compared with a cold-start model, where the initial parameters of the model were obtained from random initialization. The molecule optimization procedure is defined as inputting a molecule and returning an optimized molecule with a similar structure and better properties (**Fig. 7a**). Note that the molecule optimization task differs from the molecular de novo generation task in because it input a molecule and returns an optimized molecule, while generation tasks generally require no input and generate a set of molecules.

**Fig. 7.**
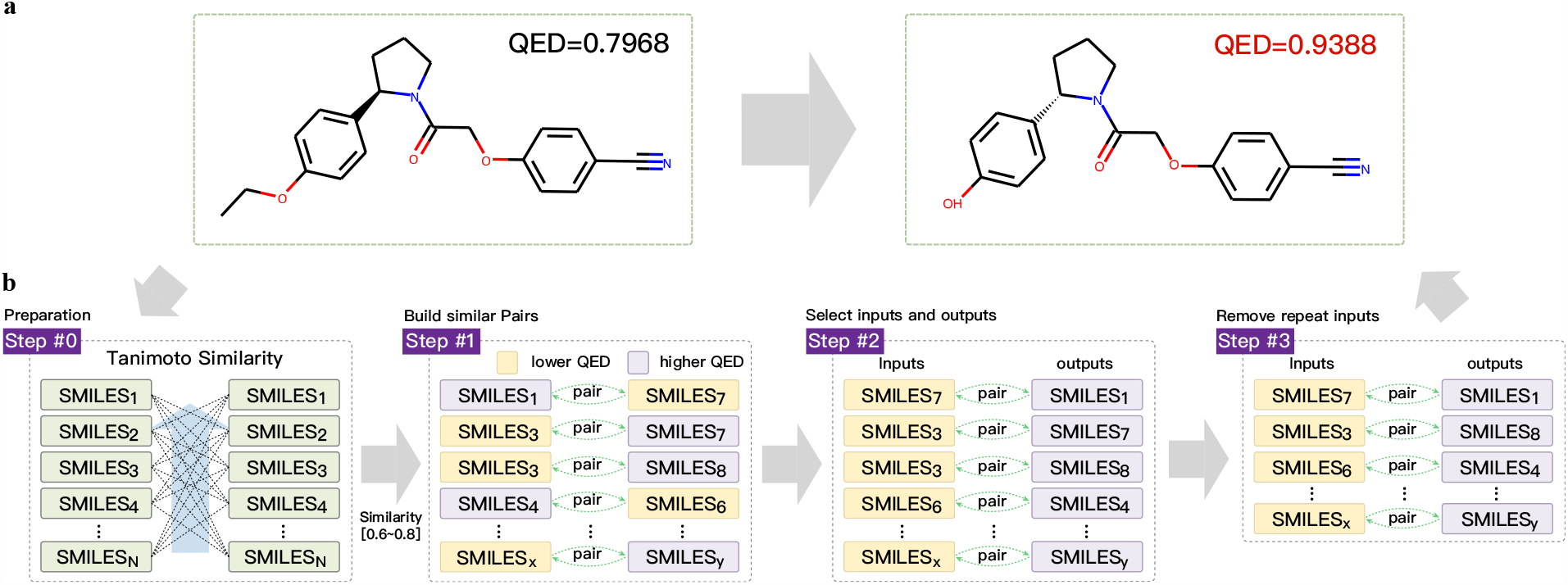
Molecule optimization by X-MOL. **a)** Similar molecules with different properties. **b)** The process of data construction for molecule optimization.

Specifically, we defined a data transformation strategy for molecule optimization with the QED as the optimization goal; i.e., we aimed to optimize a given molecule with an improved QED value. For a labelled dataset containing *n* molecules with their original QED values, we first calculated the similarity between all the molecules within the dataset, resulting in *n*(n-1)/*2 molecular pairs with their calculated similarities (**Fig. 7b**). The Tanimoto similarity was applied here, which was calculated by RDKit. First, we set a similarity range of [0.7 ± 0.1] based on the distribution of the internal similarity of the training data, and the molecular pairs that had similarities in this range were selected. Then, in each molecule pair, the molecule with a lower QED value was taken as input, and the molecule, with a higher QED value, was taken as output. To reduce redundancy, for pairs with identical input molecules, we retained only the pair with the largest difference in the QED values between the two molecules. Finally, these molecule pairs were taken as the training data to fine-tune X-MOL to obtain a molecule optimization model. In the evaluation of the molecule optimization task, we focussed on the novelty and validity rather than the uniqueness since we aimed to modify the given molecule to obtain a novel and valid molecule instead of re-sampling from the training sets. In addition, we calculated (1) the similarity between the input and output molecules and (2) the improvement of goal value compared between output molecules and input molecules. The larger these two measurements are, the better the optimization results.

### 4.3 Visualization and interpretation of the fine-tuning of X-MOL on diverse molecular analysis tasks The mechanism of attention for molecule representation

In the attention mechanism of molecule representation, the relation between two related characters was highlighted, while the irrelevant characters among them were skipped, as with a character that pays attention to the characters that matter and ignores the others. For example, the two ***c1*** instances in ***O=C(O)c1cccc(P(=O)(O)O)c1*** should pay more attention to each other than the other characters. When calculating the attention for each character, the feature vector was first mapped to ***Q***_***i***_ (the query of character *i*), ***K***_***i***_ (the key of character *i*) and ***V***_***i***_ (the value of character *i*), and the training process of the model was to identify an appropriate mapping for ***Q, K*** and ***V***. The attention ***Z***_***i***_ of character *i* was calculated according to **equation (1)**.

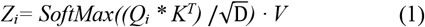

In equation (1), ***K*** is a matrix representing all the keys of the characters. ***D*** is the dimension of the feature vector. Softmax was applied to the dot product ***Q***_***i***_ ***\* K***^***T***^ to determine the weight of attention that character *i* paid to all the characters, including itself. Finally, a weighted sum of all the values of the characters was calculated as the new attention ***Z***_***i***_ of character *i*.

Generally, multiple relations exist between characters; therefore, a multi-head mechanism is adopted in practice. ***Q***/***K***/***V*** differs among different heads, and this equips the model with the ability to focus on different relations. In our study, X-MOL used 12 heads to capture at most 12 different relations between the characters.

#### The visualization of attention

The visualization of attention is actually the visualization of the attention weights. The attention weights, in the form of a two-dimensional matrix, represent the level of attention that each character pays to all the characters, including itself, in the SMILES. The attention is eventually represented in a heat map, where the brighter areas represent a stronger level of attention. Therefore, by visualizing the attention weights as a heat map, we can intuitively interpret how the model learns and understands the SMILES representation.

## Acknowledgement

This work was supported by the National Key Research and Development Program of China (Grant No. 2017YFC0908500, No. 2016YFC1303205), National Natural Science Foundation of China (Grant No. 31970638, 61572361), Shanghai Natural Science Foundation Program (Grant No. 17ZR1449400), Shanghai Artificial Intelligence Technology Standard Project (Grant No. 19DZ2200900) and Fundamental Research Funds for the Central Universities.

## Author’s contributions

Q.L. and Y.K.L. conceived the study. D.Y.X. and H.Z. implemented X-MOL. D.L.X., Y.K.G. and C.G.H. performed data collections and related analysis. Y.S., H.T. and W.H. provided valuable discussions and assistances throughout the study. Q.L., D.Y.X and Y.K.L wrote the manuscript with assistance from other authors.

